# Epithelial monolayer development and tight junction assembly on nanopillar arrays

**DOI:** 10.1101/2022.03.09.483692

**Authors:** Jose Yeste, Xavi Illa, Nitesh Shashikanth, Anton Guimerà-Brunet, Rosa Villa, Jerrold R Turner

## Abstract

Nanostructured materials provide an outstanding opportunity to both stimulate and measure cellular processes. In the context of tight junctions, it was previously reported that transient application of a nanotopographic surface over the apical brush border membrane of epithelial monolayers triggers redistribution of ZO-1, claudins, and F-actin that increases paracellular macromolecular flux. In excitable tissues, nanomaterials have been used to apply and measure electrical signals, such action potentials. As a first step towards translating these technologies for use in analysis of epithelial function, we sought to culture monolayers composed of transporting epithelia over nanopillar arrays without perturbing cellular structure or function. Madin-Darby Canine kidney I (MDCK I) cells were cultured on collagen-coated silicon chips with ∼1 μm diameter nanopillar arrays. Fluorescence and scanning electron microscopy were used to assess the impact of height on nanopillar-epithelial interactions. Monolayers formed over and were largely unaffected by short nanopillars. These nanopillars were located beneath basal epithelial surfaces and were not preferentially located within lateral intercellular spaces or beneath ZO-1-containing junctions. In contrast, tall nanopillars that exceeded cell height disrupted MDCK I monolayer growth. Cells interacted with, encircled, and extended cytoplasm over the top of tall nanopillars, and dense ZO-1 and F-actin accumulations occasionally surrounded apical membranes adjacent to nanopillars. Finally, when grown over arrays composed of nanopillars 1 – 2 μm shorter than cells, MDCK I frequently grew between nanopillars. As a result, nanopillars were more commonly present within lateral intercellular spaces beneath junctions. Apical complex structure was intact, as assessed by fluorescence microscopy of ZO-1, occludin, claudin-2, F-actin, and E-cadherin. Apical microvilli were also unaffected. We therefore show that conditions can be defined to allow growth of mature, correctly assembled epithelial monolayers with nanopillars localized to lateral intercellular spaces. This sets the stage for application of nanotechnologies for perturbation and analysis of epithelial biology.

## INTRODUCTION

The ability of skin and gut epithelial barriers to separate internal and external environments is essential for life. At these sites, the paracellular space between adjacent cells is sealed by tight junctions. Recent studies have dramatically changed our perception by showing that tight junctions are remarkably active at a molecular level.^1-4^ The first tight junction protein identified, ZO-1,^5^ promotes recruitment of transmembrane claudin proteins to the nascent tight junction.^6^ Although ZO-1 knockout in mice results in embryonic death, tissue-specific deletion within intestinal epithelia has only a mild effect on barrier function and homeostasis in the absence of injury.^7^ Nevertheless, mice lacking intestinal epithelial ZO-1 are unable to effectively repair the intestinal epithelial barrier following injury. This reflects to essential contributions of ZO-1 to epithelial proliferation and structure.^7^ These functions depend on ZO-1 scaffolding functions^6, 8-11^ that direct tight junction assembly and dynamics by interactions with other tight junction proteins as well as the actomyosin cytoskeleton.^10-19^

Most recently, several groups have shown that ZO-1 undergoes phase separation to effect tight junction assembly and epithelial migration, i.e., mechanotransduction.^3, 20-23^ How ZO-1 undergoes phase transition, the intercellular signaling events that drive condensation, and downstream consequences of this process remain poorly understood. The recent observation that application of a nanostructured film to the apical surface of intestinal epithelial monolayers could trigger tight junction remodeling, ZO-1 phase separation, and formation of cytosolic junction-like complexes may provide some clues.^4^ This work also demonstrates the potential of nanotechnology in analyses of epithelial function.

Nanostructured materials have been used to both stimulate and measure cellular processes in a variety of non-epithelial cell types. Specific examples include the use of nanostructured surfaces for intracellular delivery and extraction of diverse cargos, intracellular electrical stimulation and sensing, probe-based biochemical sensing, label-free sensing, and biomechanical stimulation.^24^ To date, nanotechnology has been applied to epithelia in order to modify growth and differentiation, regulate morphology, and stimulate cellular responses.^25-27^ For example, epithelial culture on surfaces containing dense arrays of short, less than 2 μm, nanopillars or linear grooves have been shown to markedly affect epithelial morphology and adhesion. However, nanotechnology has not been applied to analysis of transcellular and paracellular transport, which are central to epithelial function. Although nanotechnology has enormous potential as a tool with which to probe epithelial biology, a major obstacle is the absence of surfaces and growth conditions that allow epithelial growth on nanostructured surfaces without perturbing cellular structure or function.

Here, we explored dimensions of nanopillar surfaces in a systematic manner in order to determine surfaces and growth conditions that allow epithelial polarization with lateral membranes apposed against nanopillars sides and apical junctional complexes formed over nanopillar tips. We found that widely-separated, high aspect ratio nanopillars that are 1 – 2 μm shorter than epithelial cells allowed development of mature monolayers in which nanopillars were frequently present within lateral intercellular spaces, i.e., beneath junctions. Importantly, this did not appear to affect epithelial structure or function. The successional definition of these conditions is an essential first step in the application of nanotechnologies to analyses of epithelial biology.

## MATERIALS AND METHODS

### Chip and nanopillar synthesis

Patterns of photoresist dots on the 4” wafers were performed using standard photolithography process and a mask aligner Karl Süss MA6. Photoresist AZ ECI 3012 (Microchemicals GmbH) was spin coated on the wafers at 4000 rpm obtaining a 1.2 μm thick photoresist layer. Deep reactive ion etching process was conducted on an ALCATEL 601 E and numbers of etching cycles were selected to obtain a range of nanopillar heights, i.e., 4.5, 5.7, 6.5, and 8.6 μm. After the etching process, a dry thermal oxidation of the wafers was performed in an ASM LB45 furnace at 1100 ºC; thickness of the growth SiO_2_ was 0.5 μm. This SiO_2_ was then wet etched from the surface of the wafer to reduce the diameter of the nanopillars. Wafers were then thermally oxidized again to generate a thin layer of SiO_2_ of 50 nm in thickness. As a final step to produce individual chips, wafers were cut into 10×10 individual chips. Every chip included an array of nanopillars of ∼1 μm in diameter and a height of 4.5, 5.7, 6.5, or 8.6 μm. As orientation marks on the chips that allow localization and identification of nanopillars. These large structures were forming a grid of 0.8 × 0.8 mm squares on the chip.

### Cell lines and cell culture

Madin-Darby canine kidney (MDCK) I cells expressing mCherry-ZO-1 and tet-on inducible mEGFP-claudin-2 were used for all experiments. For mEGFP-claudin-2 eexpression was induced by supplementing media with 50 ng/ml of doxycycline over the last 2 days of culture. MDCK I cells were maintained in low glucose (1 g/l) DMEM supplemented with 10% fetal bovine serum (FBS), 15 mM HEPES, and G418 (250 μg/ml) and hygromycin B (50 μg/ml).

For culture, chips were transferred to individual wells of a 24 multiwell plate and washed with 70% ethanol for sterilization. Ethanol was removed and 300 μl collagen I in 70% ethanol (20 μg/ml) was dropped onto each chip. After overnight evaporation at room temperature, cell suspension (500 μl) was plated onto the chips using 500 μL of media with cells in suspension (∼500 × 10^3^ cells/ml). Cells were incubated at 37 °C and 5 % of CO_2_ and maintained for 6 days refreshing media every day.

Cell suspension (166 μl, (∼40 × 10^3^ cells/ml) was added to the apical chamber of each Transwell 3470 (Corning) and 1 ml of growth media without selective agents was placed into the basolateral chamber. Media was replenished on days 2, 4, and 6.

### Fluorescent staining and imaging

Cells on Transwells or chips were fixed in periodate-lysine-1% paraformaldehyde^28^ for 20 min at room temperature. After washing with PBS and blocking with 50 mM NH_4_Cl prepared in PBS, cells were permeabilized with Triton X-100 (0.1%) in PBS, and blocked with PBS containing 10% donkey serum. Primary antibodies diluted in blocking buffer were added. After incubation, cells were washed with PBS containing 2% donkey serum and secondary antibodies were added. After incubation and washes, Transwell filter membranes were excised. Membranes or chips were mounted under coverslips using Vectashield plus (Vector Labs). Antibodies used were monoclonal mouse anti-E cadherin clone M168 (Abnova), monoclonal rat anti-occludin clone 6B8A3, monoclonal rat anti-ZO-1 clone, and donkey anti-mouse or anti-rabbit Ig conjugated to Alexafluor 647 (Jackson Immunoresearch). For some cells, antibodies were omitted and either Alexafluor 647 phalloidin (Life Technologies) or CF640R wheat germ agglutinin (Biotium). Hoechst 33342 (0.5 μg/ml) was routinely included with secondary antibodies in order to detect nuclei. EGFP-claudin-2 and mCherry-ZO-1 were detected based on the fluoroprotein tag.

Fluorescence micrographs were collected using a DM400 (Leica) epifluorescence microscope equipped with HC PLAN APO 20x/ NA 0.70 and HCX PL APO 100x/ NA 1.35 objectives, AT350/50x, ET470/40x, ET560/40x, and ET640/30x filter cubes (Chroma), and X-cite light source (Excelitas), and a HQ2 CCD cameras (Roper) controlled by Metamorph 7 (Molecular Devices). Image stacks were collected at 1 μm or 0.2 μm intervals for low and high magnification objectives, respectively. Postacquisition processing used Metamorph and ImageJ.

### Focused ion beam-scanning electron microscopy

Combined focused ion beam-scanning electron microscopy (FIB-SEM) was used for obtaining cross section high-resolution images of the nanopillars and the cells. Sample preparation was performed embedding cells on chips with resin. Fixation for FIB-SEM was conducted in two steps. First, cells were fixed using 3 % paraformaldehyde in PBS for 20 min at room temperature; subsequently, cells were fixed in a glutaraldehyde fixative overnight at 4ºC (2.5% glutaraldehyde, 146 mM sucrose in 0.1M sodium cacodylate). Cells were washed cacodylate buffer 3 times and incubated in 1 % osmium tetroxide (OsO4) in cacodylate buffer for 1 h in ice. Then, samples were washed with 2% uranyl acetate and incubated with fresh 2% uranyl acetate at 4 ºC in dark. Dehydration of the samples were carried out with two incubations of 10 min for each dilution of ethanol (50, 75, 90, 95 % ethanol) and three final incubations using 100 % ethanol. Samples were incubated with Embed 812 resin / ethanol (1:1 v/v) at room temperature overnight followed by three incubations with fresh Embed 812 resin (2-3 h each) and a final incubation at 60ºC for 2 days for curing. To identify nanopillar locations without polishing, the silicon chip was separated from the resin in a process that left the nanopillars embedded in the plastic. This process consisted in repeated cycles of adding the sample in liquid nitrogen and transferring it to a beaker with water. The embedded samples then consisted of cells and the nanopillars on the surface of the block. Preparation for SEM was conducted mounting the samples in a SEM slab, painting the lateral of the sample with silver paint, and sputtering 10 nm of platinum/palladium. Samples were imaged using a Zeiss 1560XB Cross Beam at a magnitude 7000 and backscattered detector. Cross sections were obtained with slices every 100 nm.

### Statistics

Data are expressed as the mean ± SD or mean ± interquartile bar. Statistical analyses were performed using ANOVA in GraphPad Prism 9. Results were considered significant at p < 0.05.

## RESULTS

### Nanopillar chip development

Si/SiO_2_ chips with arrays of nanopillars intended to localize between cells were produced. The MDCK I epithelial cells used here are flat to cuboidal, with cross sectional diameters of ∼20 μm and heights of up to ∼8 μm, when grown as confluent monolayers on semipermeable Transwell supports (Fig. 1A). We reasoned that junctions would not assemble over nanopillars that are taller than the cells. Conversely, very short nanopillars would not be expected to reach the subjunctional space. We therefore anticipated that nanopillars would have to be between 5 μm and 10 μm tall. To accomplish this while using small nanopillars unlikely to affect cell behavior required use of high-aspect ratio nanopillars.

**Figure 1.**
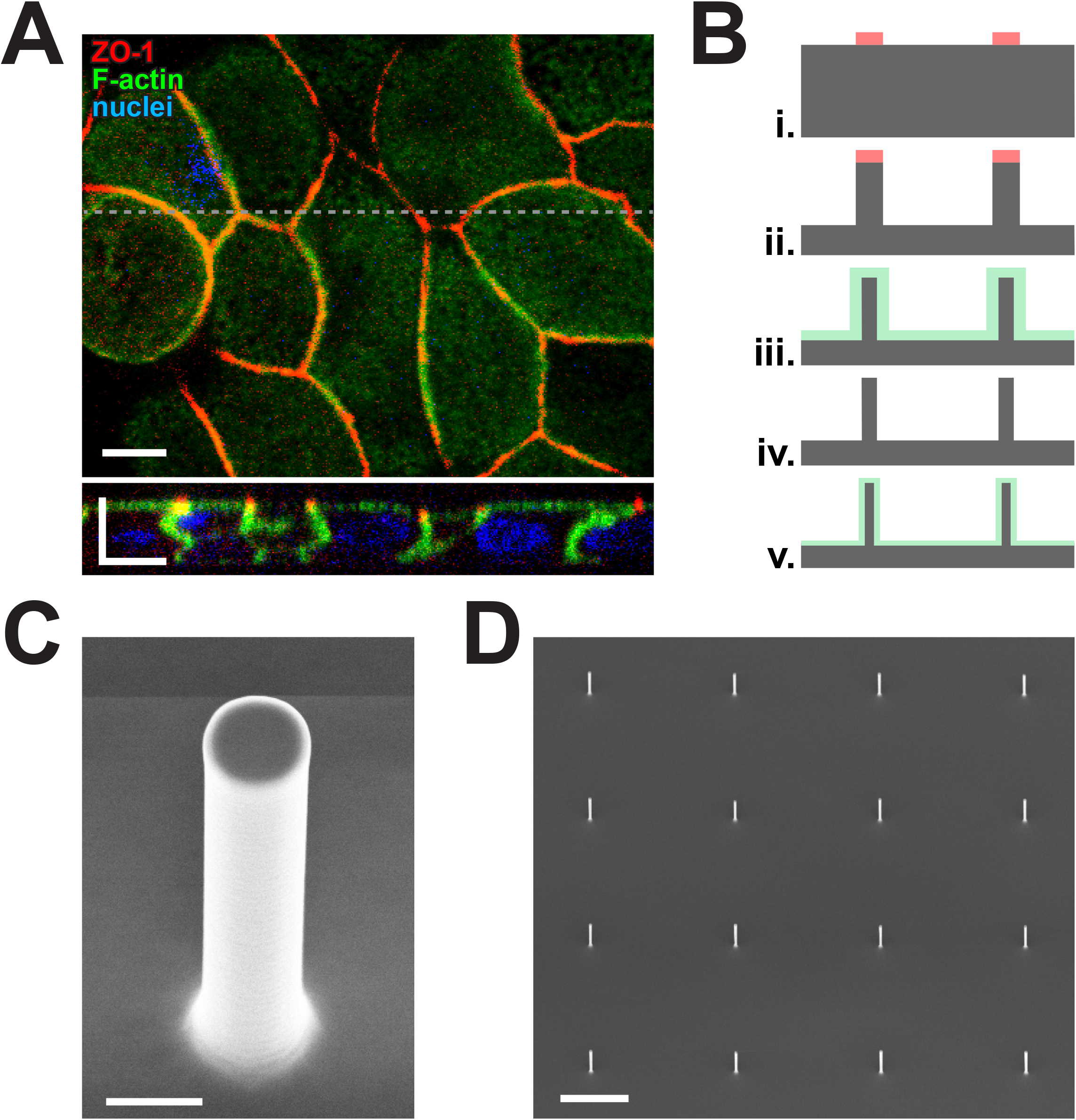
MDCK I cell morphology and nanopillar chip design. ***A*.** MDCK I monolayers expressing mCherry-ZO-1 were stained for F-actin and nuclei. Images (*xy*-plane and *xz*-plane) are shown. The dashed line in the *xy* image indicates the position from which the *xz* image was obtained. The *xy* image is taken at the plane of the tight junction. Bars = 10 μm. ***B***. Schematic of chip synthesis. i) Photoresist dots (pink) patterned on silicon wafer (grey). ii) Deep reactive ion etching leaves Si nanopillars with photoresist tips. iii) Thermal oxidization to create a 0.5 μm thick SiO_2_ layer (green) over the surface. iv) Wet etching removal of the SiO_2_ layer. v) Thermal oxidization to create a 50 nm SiO_2_ layer (green) over the surface. ***C***. Scanning electron micrograph of a nanopillar. Bar = 1 μm. ***D***. Low magnification scanning electron micrograph of a nanopillar array. Bar = 10 μm.

Chips including nanopillars were fabricated on using 4” Si wafers. Tall and thin structures were created by combining processes of dry etching, thermal oxidation, and wet etching (Fig. 1B). We first patterned photoresist dots (1.4–1.8 μm in diameter) on the wafer using a photolithography process and used those dots as a soft mask to etch the silicon in a deep reactive ion etching process. Preliminary nanopillars had an aspect ratio of ∼2.5–6. To increase this value, nanopillars were thermally oxidized obtaining a 0.5 μm thick SiO_2_ layer over the structures. The SiO_2_ material was then selectively wet etched obtaining in order to create Si nanopillars with aspect ratios of at least 10. In order to generate nanopillars similar in height to MDCK I cells, chips were generated with 1 μm diameter nanopillars that were 4.5, 5.7, 6.5, and 8.6 μm tall (Fig. 1C). The spacing between nanopillars was set to 20 μm in order to allow cells to grow onto the nanopillar surface with little interference (Fig. 1D). As a final step, wafers were thermally oxidized one last time to create a thin, glass-like 50 nm SiO_2_ layer, as many epithelial cells, including MDCK I cells, grow well on glass surfaces, either directly or after modification. Finally, wafers were diced into 10 × 10 mm chips for use.

### Optimizing cell growth

Density at plating is well-recognized to impact epithelial monolayer development. When grown on semipermeable supports, i.e., Transwells, MDCK I cells are typically plated at a density of 20,000 cells/cm^2^. To assess the impact of density when cells are grown on chips with a SiO_2_ surface, MDCK I cells were plated at concentrations from 3,000 to 100,000 cells/cm^2^ and allowed to assemble into monolayers over 6 days of growth. Although cells became confluent when plated at low densities, they were only ∼5 μm tall, significant way shorter than cells grown on Transwells (Fig. 2A). This corresponded to significant greater cell areas (Fig. 2B). Thus, MDCK I cells had a flat, squamoid shape when plated at low densities on the SiO_2_ chip surface. Greater plating densities resulted in progressive increases in MDCK I cell height. Plating at 50,0000 - 100,000 cells/cm^2^ resulted in monolayers with heights of ∼8 μm, similar to that of monolayers grown on Transwells. However, the highest density of plating (100,000 cells/cm^2^) led to much more uniform cell heights than lower concentrations. Conversely, cell area decreased as plating densities increased. Cells plated at 100,000 cells/cm^2^ had surface areas comparable to those of Transwell-grown monolayers. Similar to cell heights, the range of cell areas was more limited at the highest plating density.

**Figure 2.**
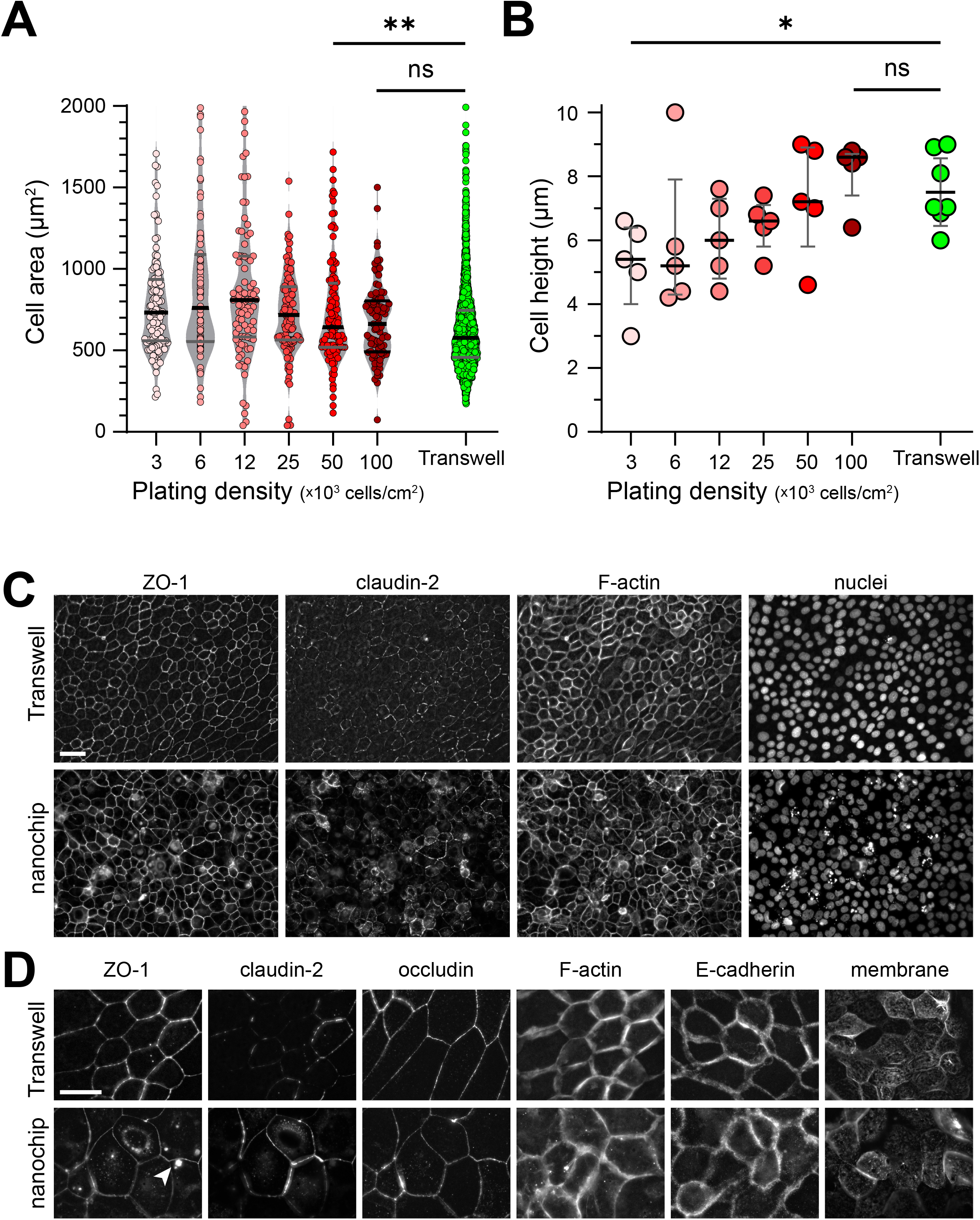
Morphology of cells grown on Transwells and chips. ***A*.** Low magnification views of mCherry-ZO-1, EGFP-claudin-2, F-actin, and nuclei. Morphologies are similar with the exception of scattered apoptotic bodies within monolayers cultured on chips. Bar = 50 μm. ***B***. High magnification views of mCherry-ZO-1, EGFP-claudin-2, occludin, F-actin, E-cadherin, and wheat germ agglutinin-labeled cell membranes. Note the cytoplasmic ZO-1 aggregates in some cells. These did not colocalize with nanopillars and represent an overexpression artifact. Bar = 25 μm. ***C***. Cell area was measured for cells grown on Transwells or Si/SiO_2_ chips at the indicated plating densities. Mean (black) and interquartile (grey) lines. ***D***. Cell height from the growth surface to location of mCherry-ZO-1 was measured for cells grown under conditions indicated. Mean ± SD.

In order to assess monolayer integrity, the morphology of MDCK I cells plated at 100,000 cells/cm^2^ was compared to that of Transwell-grown cells. In both cases the uniform distribution of cells could be readily recognized by stain for the tight junction protein ZO-1 (Fig. 2C). Moreover, stably transfected MDCK I lines expressed the transgene, EGFP-claudin-2, equally well under both growth conditions and readily localized claudin-2 at tight junctions (Fig. 2C,D). Similarly, microfilament, i.e., F-actin, architecture was unperturbed by growth on chips and was indistinguishable from that of MDCK I grown on Transwells. The only difference between cells grown on the two surfaces was detected by nuclear (DNA) stain. Although relatively few in number, brightly-stained nuclear fragments, typical of apoptotic debris, were present overlying chip-grown monolayers.

To further assess intercellular junctions, the distributions of ZO-1, EGFP-claudin-2, occludin, F-actin, E-cadherin were assessed at greater magnification (Fig. 2D). Morphology was were also assessed using fluorescent-tagged wheat germ agglutinin, which binds to N-acetyl-D-glucosamine and sialic acid and effectively labels cell membranes (Fig. 2D). The tight junction proteins co-localized at intercellular junctions. Maximum projections of F-actin, E-cadherin, wheat germ agglutinin labeling show that both surface architecture and lateral membrane structures were unperturbed by growth on chips.

### Effect of nanopillar height

MDCK I cells expressing EGFP-claudin-2 grown on chips with 4.5, 5.7, 6.5, and 8.6 μm tall nanopillars. In all cases, the pattern of growth was such that each 20 μm × 20 μm square defined by 4 nanopillars (Fig. 3A). This resulted more than ∼40% of nanopillars being situated within or very close to the lateral intercellular space. It was not difficult to find nanopillars beneath the tight junctions, as defined by EGFP-claudin-2. The images show one example of a nanopillar tip directly under the EGFP-claudin-2-labeled tight junction (Fig. 3A). Monolayers were composed of cuboidal cells placed primarily between nanopillars; cell area and height remained to those of Transwell-grown monolayers.

**Figure 3.**
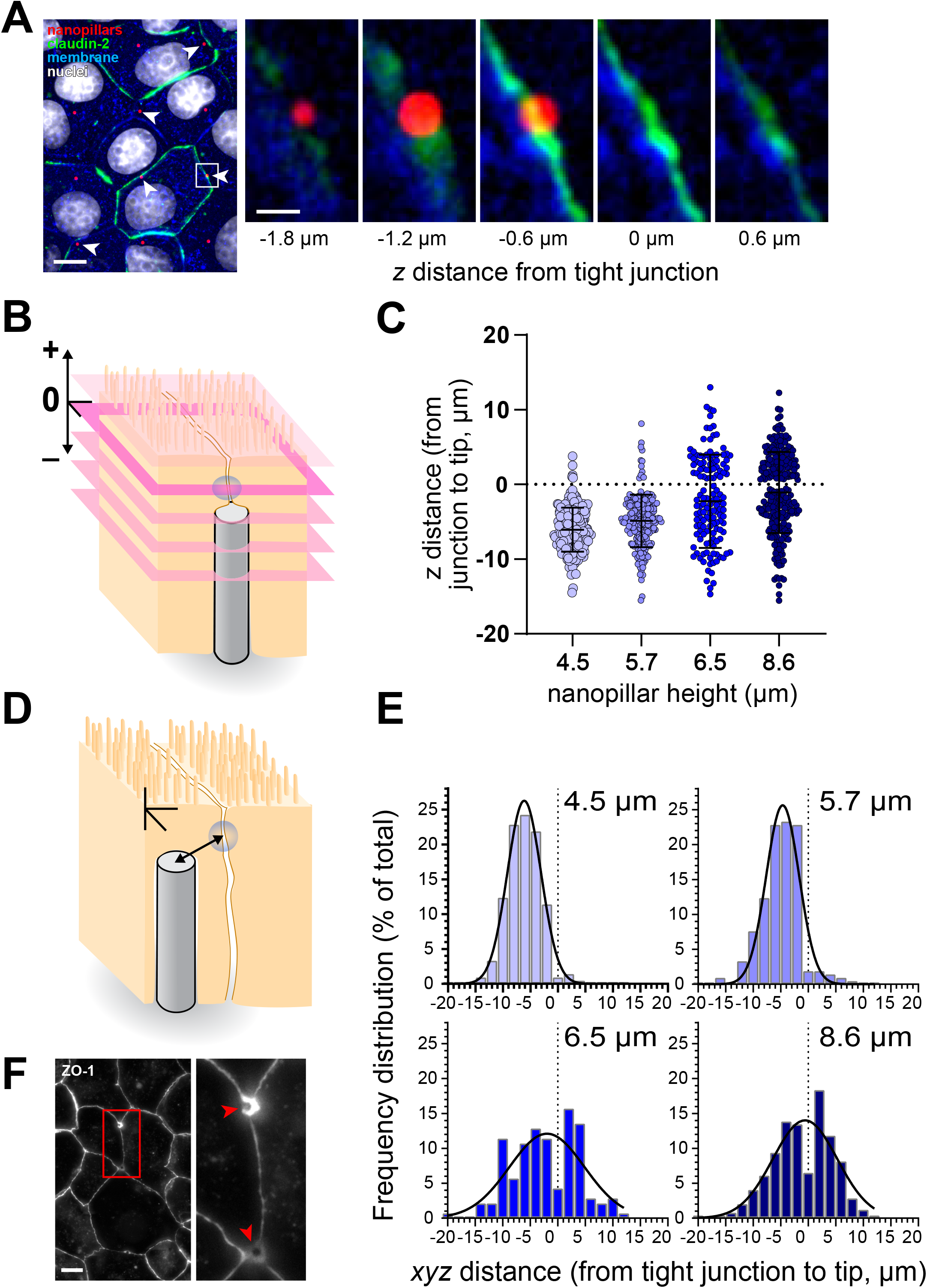
Nanopillar positioning relative to tight junctions. ***A*.** Low-power view of nanopillars (red), EGFP-claudin-2 (green), wheat germ agglutinin staining (blue) and nuclei (white). Many nanopillars are located at or immediately adjacent to claudin-2-labeled tight junctions (arrows). The insets show the relationship between the nanopillar and the junction in the *z*-axis. Note how the nanopillar tip is just below the top of the EGFP-claudin-2 band defining the tight junction. *z*-plane distance from the tight junction is indicated beneath each image, with negative values indicating positions below the tight junction. Bars = 10 μm, 1 μm (insets). ***B***. The vertical (*z*-plane) distance from the tip of each nanopillar (grey) to the *xy*-plane (pink) of the mCherry-ZO-1-labeled tight junction (blue) was determined. Positive values indicate nanopillar tips above the tight junction plane, negative values indicate tips below the tight junction plane. ***C***. *z*-plane distance as a function of nanopillar placement relative to the tight junction, determined as in (B). ***D***. The 3-dimensional (*x. y, z*-planes) distance from each nanopillar tip to the closest mCherry-ZO-1-labeled tight junction (blue) was determined. values indicate nanopillar tips above the tight junction plane, negative values indicate tips below the tight junction plane. **E**. 3-dimensional distance from each nanopillar tip to the tight junction. F. Particularly with 8.6 μm tall nanopillars, some sites displayed bright semi-circular mCherry-ZO-1 surrounding one side of the adjacent nanopillar (arrows).

To assess the impact of nanopillar height on the relationship between nanopillar and tight junctions, monolayers were fixed and stained with ZO-1, whose distribution in the *z*-axis is much more restricted than that of claudin-2 and occludin. The z-axis distance from the ZO-1-labeled tight junction to the tip of each nanopillar was determined (Fig. 3B). As expected, when monolayers were grown on 4.5 μm nanopillars, the nanopillar tips were nearly always located significant distances from the z-plane of the tight junction, with a mean of ∼ 7 μm (Fig. 3C). Only rare, flattened cells had nanopillars that extended above the tight junction. Increasing nanopillar height to 5.7 μm did not appreciably improve this, and most nanopillar tips were well below the tight junction (Fig. 3C). When grown on 6.5 μm tall nanopillars, the majority of tips were located within ∼3 μm of the tight junction z-plane, with a mean of less than 2 μm (Fig. 3C). However, use of taller nanopillars resulted up to 35% of tips being above the tight junction. This problem was exacerbated when monolayers were grown on 8.6 μm tall nanopillars.

Although the vertical distance between nanopillar tip and ZO-1-decorated tight junction is a critical parameter, the distance as measured in *x*-, *y*-, and *z*-planes (3 dimensions) provides important information (Fig. 3D). After growth on 4.5 μm tall nanopillars, more than 50% of nanopillar tips were greater than 5 μm from the closest tight junction. This improved slightly when nanopillars were extended to 5.7 μm (Fig. 3E).

Use of 6.5 μm and 8.6 μm nanopillars led to significantly greater numbers of tips within ∼3 μm of the tight junction. These were equally divided between nanopillar tips above or below the tight junction for 6.5 μm tall nanopillars but were skewed towards tips above junctions with the 8.6 micro nanopillars (Fig. 3E). However, the tips of some taller nanopillars became encircled by cell processes (Fig. 3F). Although this occurred with both 6.5 μm and 8.6 μm nanopillars, it was more frequent with the tallest nanopillars. Thus, the data indicate that 6.5 μm is the optimal nanopillar height for MDCK I cells.

### Nanopillar tip apposition directly beneath tight junctions

The qualitative immunofluorescence microscopy as well as quantitative morphometry suggests that nanopillars are directly beneath tight junctions, but the resolution of light microscopy, particularly in the *z*-axis, is limited. We therefore examined nanopillar chips with cultured cells by focused ion beam scanning electron microscopy (FIB-SEM) using sections oriented in the *xz*-plane (relative to the immunofluorescence). Although not all nanopillars were adjacent to junctions, serial views along the *y*-axis of a representative nanopillar show that the tight junction between two cells is almost directly above the nanopillar tip (Fig. 4). Given the limited sampling possible by FIP-SEM, the relative ease with which the nanopillar shown was identified indicate that sufficient numbers of nanopillars will be correctly placed.

**Figure 4.**
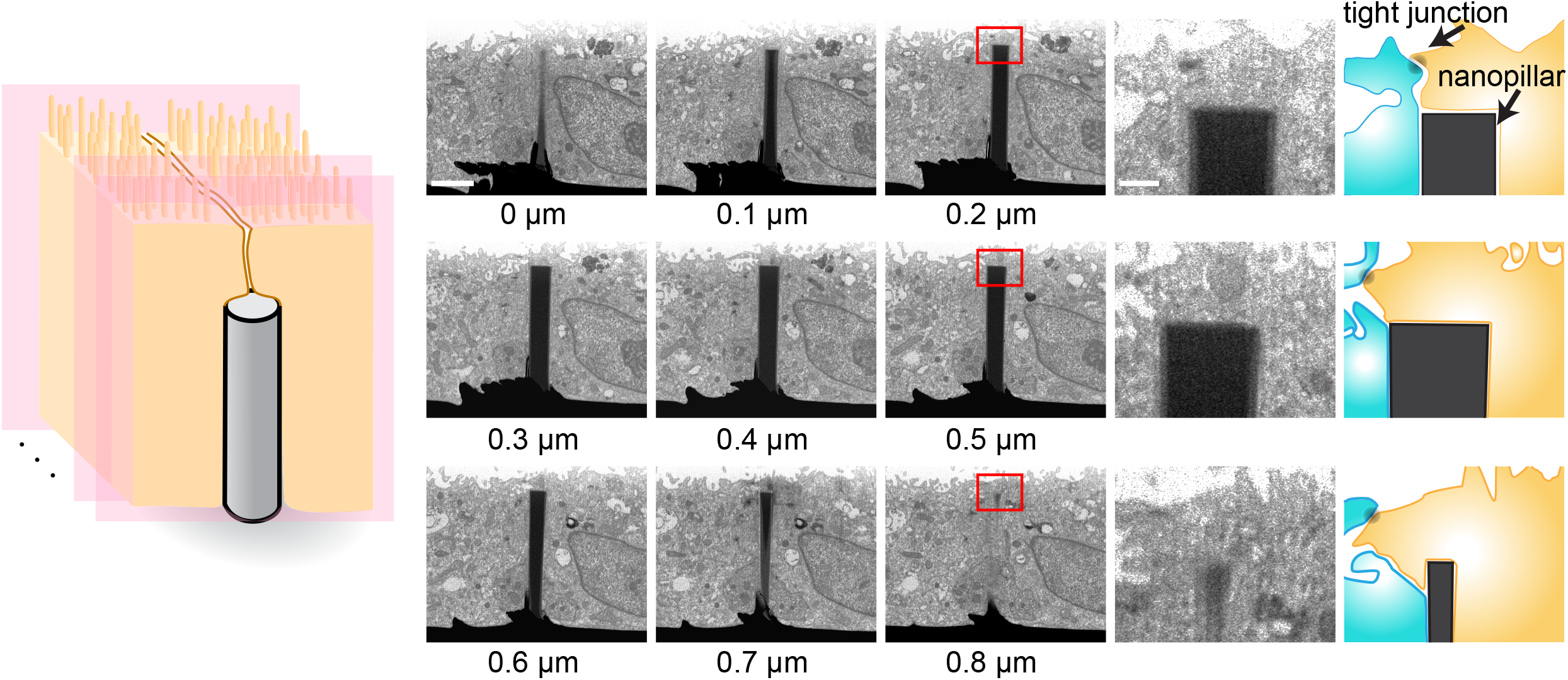
Relationship between nanopillars and epithelial cells as imaged by FIB-SEM. Successive images were collected at 0.1 μm intervals. The diagrams to the right of each inset shows the position of two cells and the tight junction relative to the nanopillar. Bars = 2.5 μm, 0.5 μm (insets).

## DISCUSSION

Epithelial surfaces are formed by a continuous layer of cells. Integrated function across the monolayer requires specialized intercellular junctions, most critically the apical junctional complex composed, from apical to basal, of tight junctions, adherens junctions, and desmosomes.^29^ These structures are highly dynamic and, therefore, susceptible to perturbation by external stimuli. Thus, based on previous experiences where nanopillars were used to probe or deliver substances to the cytoplasm, we had initial concern that it would not be possible to grow epithelial monolayers on nanopillar arrays without interfering with normal structure and function. Here, as a prerequisite to assessing function, we have developed and tested nanopillar arrays that allow cell growth. Moreover, we have identified culture conditions that allow epithelial cells form monolayers that are structurally nearly indistinguishable from those grown on semipermeable supports, which can be considered the “gold-standard.”

One important parameter to consider is cell area. When epithelial damage occurs, cells surrounding the wound flatten and extend to reform the barrier via restitution. During this process cell area increases while height decreases. It was, therefore, critical to determine conditions that allowed epithelial cells within confluent monolayers to develop with areas and heights similar to that of Transwell-grown cells. We found that this required a plating density 5-fold greater than that necessary for growth on Transwells. This is comparable to cell density density of mature MDCK I monolayers and suggests that the Si/SiO_2_ chip surface impaired the ability of cell populations to expand and migrate to form a confluent monolayer of cuboidal, rather than squamoid (flat), cells. Consistent with this we also noted a slight increase in apoptosis when cells were grown on chips relative to Transwell membranes. Nevertheless, when plated at the higher concentration monolayers formed and most squares marked by nanopillars spaced 20 μm from one another contained a single epithelial cell. It is, therefore, possible to create mature epithelial monolayers on Si/SiO_2_ nanopillar chips.

Although some nanopillars were present beneath cell bodies, rather than within lateral intercellular spaces, this did not appear to interfere with intercellular junction assembly. Some bright foci of cytoplasmic ZO-1 were occasionally present and often colocalized with nanopillars. These could represent phase separation,^2^ but the observation that they were not associated with tight junctions suggests that they may simply be secondary to overexpression, as they were only observed in cells expressing mCherry-ZO-1.

## Acknowledgments

Supported by R01 DK068271, R01 DK061931, the European Union’s Horizon 2020 research and innovation programme under the Marie Skłodowska-Curie grant agreement No 896293, and the Spanish ICTS Network MICRONANOFABS partially supported by MICINN and the ICTS ‘NANBIOSIS’, more specifically by the Micro-NanoTechnology Unit of the CIBER in Bioengineering, Biomaterials and Nanomedicine (CIBER-BBN) at the IMB-CNM. Part of the work was performed on the core facility Center for Nanoscale Systems (CNS) at Harvard University.

